# YY males of the dioecious plant *Mercurialis annua* are fully viable but produce largely infertile pollen

**DOI:** 10.1101/658708

**Authors:** Xinji Li, Paris Veltsos, Guillaume Cossard, Jörn Gerchen, John R. Pannell

## Abstract

The suppression of recombination during sex-chromosome evolution is thought to be favoured by linkage between the sex-determining locus and sexually-antagonistic loci, and leads to the degeneration of the chromosome restricted to the heterogametic sex. Despite substantial evidence for genetic degeneration at the sequence level, the phenotypic effects of the earliest stages of sex-chromosome evolution are poorly known. Here, we compare the morphology, viability and fertility between XY and YY individuals produced by crossing seed-producing males in the dioecious plant *Mercurialis annua* L., which has young sex chromosomes with limited X-Y sequence divergence. We found no significant difference in viability or vegetative morphology between XY and YY males. However, electron microscopy revealed clear differences in pollen anatomy, and YY males were significantly poorer sires in competition with their XY counterparts. Our study suggests either that the X chromosome is required for full male fertility in *M. annua*, or that male fertility is sensitive to the dosage of relevant Y-linked genes. We discuss the possibility that the maintenance of male-fertility genes on the X chromosome might have been favoured in recent population expansions, which selected for the ability of females to produce pollen in the absence of males.

## Introduction

Sex chromosomes have evolved numerous times in eukaryotes, showing a number of features that are remarkably common. They include the suppression of recombination around the sex-determining locus, and the genetic degeneration of this non-recombining region, including the loss of genes, impairment of gene function, and the accumulation of repetitive elements (Charlesworth, 1991; Charlesworth *et al.*, 2005; Bachtrog *et al.*, 2011; Ming *et al.*, 2011; Bachtrog, 2013; Abbott *et al.*, 2017). The evolutionary genetic reasons for this degeneration are reasonably well understood (Charlesworth & Charlesworth, 2000; Bachtrog, 2008). For instance, purifying selection is much less efficient in regions of low recombination because of Hill-Robertson interference between linked loci (Hill & Robertson, 1966; McVean & Charlesworth, 2000), and processes such as genetic hitchhiking (Maynard-Smith & Haigh, 1974; Rice, 1987a), background selection (Charlesworth *et al.*, 1995; Kaiser & Charlesworth, 2009) and Muller’s ratchet result in the accumulation of deleterious mutations (Muller, 1918; Gordo & Charlesworth, 2001; Bachtrog & Gordo, 2004; Engelstädter, 2008). The effects of such processes on the non-recombining region of sex chromosomes have been documented at the genomic and transcriptomic levels in a diverse range of organisms (Rice, 1996; Steinemann & Steinemann, 1998; Berlin *et al.*, 2007; Kaiser & Charlesworth, 2010), including plants (Filatov *et al.*, 2000; Liu *et al.*, 2004; Hough *et al.*, 2017). In contrast, their phenotypic effects in terms of morphology, life-history, viability and fertility have received little attention beyond potentially associated patterns of sexual dimorphism.

Although we expect non-recombining regions of the genome to be prone to degenerative processes, they may also be targets of positive selection, conferring advantages on individuals of one or both sexes (Bachtrog, 2006; Ellegren & Parsch, 2007; Zhou & Bachtrog, 2012). Indeed, alleles that confer an advantage on one sex and a disadvantage on the other (‘sexually antagonistic’, or SA, alleles) are thought to be one of the reasons for the evolution of suppressed recombination around the sex-determining locus in the first place (Charlesworth & Charlesworth, 1980; Rice, 1987b, 1992; van Doorn & Kirkpatrick, 2007). For instance, alleles that are advantageous to males but detrimental to females will increase in frequency if in tight linkage with the male-determining locus (and vice versa for female-advantageous mutations), because recombination would place them in a genetic background in which their expression is deleterious. The sexual antagonism hypothesis also provides a plausible explanation for the existence of ‘evolutionary strata’ on sex chromosomes, where regions close to the sex-determining locus, for which recombination was suppressed first, are more divergent than those further away that stopped recombining more recently (Charlesworth *et al.*, 2005; Bergero & Charlesworth, 2009). However, despite its conceptual plausibility, there is still limited empirical evidence for SA selection on sex chromosomes (Ironside, 2010). In the guppy (*Poecilia reticulate*), attractive colouration increases male siring success but would be deleterious in females that are rendered more visible to predators (Brooks, 2000). Male colouration factors are linked to the sex-determining region of the Y chromosome in guppy populations prone to high predation (Lindholm & Breden, 2002; Charlesworth, 2018), pointing to the possibility that the non-recombining region may have expanded as a result of SA selection. However, recent work has dismissed the idea of evolutionary strata on the Y chromosome in guppies because recombination is suppressed in males generally – and not just around the sex-determining locus (Bergero *et al.*, 2019).

More widely, the enrichment of genes with sex-biased expression on the sex chromosomes is also consistent with the SA hypothesis (Ellegren & Parsch, 2007; Bachtrog *et al.*, 2011; Connallon & Clark, 2013). This is because differential gene expression between the sexes points to a response to sex-specific selection (Connallon & Clark, 2010; Meisel *et al.*, 2012). In the dioecious plant *Silene latifolia*, for instance, quantitative trait loci associated with sexual dimorphism, which have the potential to resolve intralocus sexual conflicts, show sex-specific expression (Scotti & Delph, 2006; Delph *et al.*, 2010) and are enriched on the sex chromosomes (Zemp *et al.*, 2016). Similarly, in the subdioecious plant *Fragaria virginiana*, linked sex-limited QTL underlie sexually dimorphic traits (Spigler *et al.*, 2011). Importantly, sex chromosomes in which recombination has ceased recently may come to harbour sex-specific variation before accumulating deleterious mutations, with a possible net positive effect on the corresponding sex.

The detection of phenotypic effects of Y-chromosome evolution, whether negative or positive, is likely to differ between the haploid and diploid phases of the plant life cycle. In the haploid phase of the life cycle, notably during pollen-tube growth, deleterious mutations should have an immediate effect on the phenotype. Haploid expression should therefore subject genes to strong purifying selection (Bergero & Charlesworth, 2009; Chibalina & Filatov, 2011), resulting in a slower rate of Y-chromosome degeneration in plants than in animals (Chibalina & Filatov, 2011; Charlesworth, 2015; Krasovec *et al.*, 2018). Nevertheless, the gradual accumulation of deleterious Y-linked mutations might eventually become detectable in the form of reduced competitiveness of Y chromosome-bearing pollen tubes (Charlesworth, 1988; Sandler *et al.*, 2018), a phenomenon known as ‘certation’ (Smith, 1963; Lloyd, 1974). Such effects on gametophyte fitness have been invoked to explain female-biased sex ratios in the wind-pollinated dioecious plant *Rumex nivalis* (Stehlik *et al.*, 2008). In contrast to their effects on the haploid phase of the plant life cycle, recessive mutations on the Y chromosome will tend to be masked in the diploid phase as a result of the expression of functional alleles on the X, and even partially dominant mutations might become hidden by dosage compensation (Charlesworth, 1996; Wilson & Makova, 2009; Vyskot & Hobza, 2015).

Although it will often be difficult to detect the effects of Y-linked mutations on diploid individuals, they should in fact become apparent through comparisons between XY and YY males. YY males can be generated by crossing XY males that have been artificially feminized (Durand & Durand, 1984; Khryanin, 2002) or that show natural ‘leaky’ or ‘inconstant’ sex expression (Ehlers & Bataillon, 2007; Cossard and Pannell, 2019). Such manipulations have typically resulted in nonviable YY progeny in most animals with old sex chromosomes (Graves, 2006). In contrast, YY individuals are viable and fertile in animals with non-degenerate Y chromosomes, such as a variety of fish species (Yamamoto, 1963, 1975; Chevassus *et al.*, 1988; Scott *et al.*, 1989; Kavumpurath & Pandian, 1993). A notable exception to this pattern is provided by WW individuals of the androdioecious clam shrimp *Eulimnadia texana*, in which ZW and WW hermaphrodites naturally co-occur with ZZ males. Here, WW hermaphrodites are viable but have substantially lower fitness than their ZW counterparts, pointing to recessive deleterious effects of W-linked alleles (Sassaman & Weeks, 2002). In plants, YY individuals are nonviable in both *Silene latifolia* (Janoušek *et al.*, 1998; Soukupova *et al.*, 2014), which has highly divergent heteromorphic sex chromosomes (Krasovec *et al.*, 2018), and in *Carica papaya*, which has homomorphic sex chromosomes at the cytological level but shows XY divergence at the sequence level (Liu *et al.*, 2004; Yu *et al.*, 2008b). Viable YY males have been reported in *Asparagus officinalis* (Harkess *et al.*, 2017), *Spinacia oleracea* (Yamamoto *et al.*, 2014; Wadlington & Ming, 2018), *Cannabis sativa* (Peil *et al.*, 2003), *Phoenix dactylifera* and *Actinidia chinensis* (reviewed in Ming *et al.*, 2011). Yet all these records provide limited details about the performance of YY individuals, and we are aware of no phenotypic comparisons between XY and YY individuals.

Here, we investigate the phenotypic consequences of early sex-chromosome evolution in adult males of the wind-pollinated dioecious plant *Mercurialis annua* L. (Euphorbiaceae). The *M. annua* species complex includes monoecious and androdioecious polyploid lineages, but diploid populations (which are widespread across eastern, central and western Europe) are exclusively dioecious (Obbard *et al.*, 2006). Dioecy is ancestral in the genus *Mercurialis* (Krähenbühl *et al.*, 2002), where all the dioecious species display common sexual dimorphism in their inflorescences, with male flowers developing on long peduncles held above the plant canopy and female flowers usually placed on much shorter pedicels in the leaf axils (Pannell, 1997; Buggs & Pannell, 2007). *M. annua* and its dioecious or androdioecious annual relatives share the same XY sex-determination system (Russell & Pannell, 2015). Veltsos *et al.* (2018, 2019) estimated that recombination has been suppressed over one third the length of the *M. annua* Y chromosome, a region measuring about 15 Mb with approximately 500 genes and estimated to be younger than 1 million years. About half of the approximately 30 genes that show male-specific expression in *M. annua* occur in the non-recombining region of the Y, and the possession of the pedunculate male inflorescence, a likely sexually antagonistic trait (Santos del Blanco *et al.*, 2018), is also Y-linked (Russell & Pannell, 2015; Veltsos *et al.*, 2018). Consistent with its relative youth, the Y chromosome shows signs of only mild sequence degeneration, with pseudogenisation of only a single Y-linked allele (as a result of a premature stop codon) and a modest excess in the ratio of non-synonymous to synonymous mutations compared to autosomal genes (Veltsos *et al.*, 2019).

Because the *M. annua* Y chromosome shows some signs of degeneration, we might expect YY individuals that lack an X complement to have reduced viability and/or fertility. They might also suffer fitness consequences from a dosage imbalance, since the YY genotype is rarely tested by selection. Alternatively, multiple features of the sex-linked loci, such as an enrichment in female-biased genes and the elevated sex-biased expression in male-biased genes, point to the possibility of ongoing sexually antagonistic selection shaping the Y chromosome (Veltsos *et al.*, 2019). We might therefore also expect YY individuals to have a ‘super-male’ phenotype, if the Y-linked alleles are not completely dominant. To evaluate the phenotypic consequences of the recently evolved Y chromosome in *M. annua*, we compared the phenotypes of typical XY males with experimentally induced YY males for a range of vegetative and reproductive traits. We found limited phenotypic differences in the vegetative phenotypes or in viability and vigour between XY and YY males. In contrast, the pollen produced by YY males showed signs of partial sterility, which was confirmed by the ratio of the number of XY and YY brothers to that of their XX sisters produced under open pollination. To our knowledge, our results represent the first account of the phenotypic effects of divergence between X and Y chromosomes of a plant species.

## Materials and Methods

### Generation of YY males in *Mercurialis annua*

Two approaches were taken to generate YY males in *M. annua*. In the ‘hormone’ experiment, we germinated diploid *M. annua* seeds and kept 38 male seedlings in a growth chamber. Seedlings were sprayed with feminising cytokinin solution (6-Benzylaminopurine, 2 μM) once a day, following a protocol modified from Louis & Durand (1978), which led to the production of a large number of pistillate flowers in the male inflorescences. We allowed the modified males to pollinate one another, and then bulk-harvested the seeds. In total, we collected 443 seeds from these crosses.

In the ‘pruning’ experiment, we stimulated female-flower production through severe pruning on 2,000 diploid *M. annua* males grown in a common garden, prompted by the observation reported by Kuhn (1939) that such pruning elicits increased leakiness in sex expression. After ten weeks of growth under open pollination, we harvested a total of 496 seeds from the pruned males.

### Identification and characterisation of YY versus XY males

All the seeds collected from the male crosses of both experiments were germinated and raised to maturity at the University of Lausanne. Seeds from the hormone experiment were grown in an outdoor field in 2015, and seeds from the pruning experiment were grown under glasshouse conditions in 2017. When the plants began flowering, we recorded the numbers of male and female seedlings among the progeny. In addition, a restriction enzyme-based assay was developed based on X- and Y-specific SNPs to distinguish between XY and YY males (see Supporting Information, Methods S1 & Figure S5). Briefly, a sex-linked PCR product was digested with two restriction enzymes (*BceAI* and *Rsel*) which cut only the X- or Y-linked sequence, thus allowing individuals with or without an X to be distinguished. The Y-specific SCAR marker (Khadka *et al.*, 2002) was used as a positive control for PCR amplification.

The following traits were measured on all XY and YY males: plant height, male flower biomass, peduncle biomass and total biomass. We also calculated the biomass/height ratio and the relative male reproductive allocation (MRA) as the proportion of total biomass allocated to reproductive organs, i.e., the biomass of flowers and peduncles divided by the total reproductive and vegetative biomass.

We used linear mixed-effect models to compare phenotypic traits between XY and YY individuals using the lme4 library (version 1.1-17; Bates *et al*., 2015) in R v3.5.1. We conducted principal component analysis (PCA) to eliminate covariance between traits and applied linear mixed-effect models to the first four principal components of normalised independent phenotypic measurements. All data analysis was performed using R v3.5.1 (R Development Core Team, 2008) and the summary tables from the models were generated using the R package sjPlot v2.4 (Lüdecke, 2017).

### Assessment of the relative viability of YY males

We assessed the relative viability of YY and XY males by comparing the ratio of XY and YY progeny from the crosses among feminised males, with the ratio expected from random mating and equal survival of progeny genotypes (see scheme in Figure 1a). Given the known XY sex determination in *M. annua*, crosses between parental XY males should result in a 1:2:1 ratio of XX female to XY male to YY male zygotes. Under the null hypothesis of complete YY viability, we thus expect 75% of the F_1_ progeny to be males (p1 in Figure 1a); in contrast, we expect 67% males under the assumption of complete YY nonviability (p2 in Figure 1a). An intermediate value suggests partial YY inviability. We assessed the proportion of male progeny for significant deviations from expected values using Chi-squared and likelihood-ratio tests (Etz, 2018).

**Figure 1.**
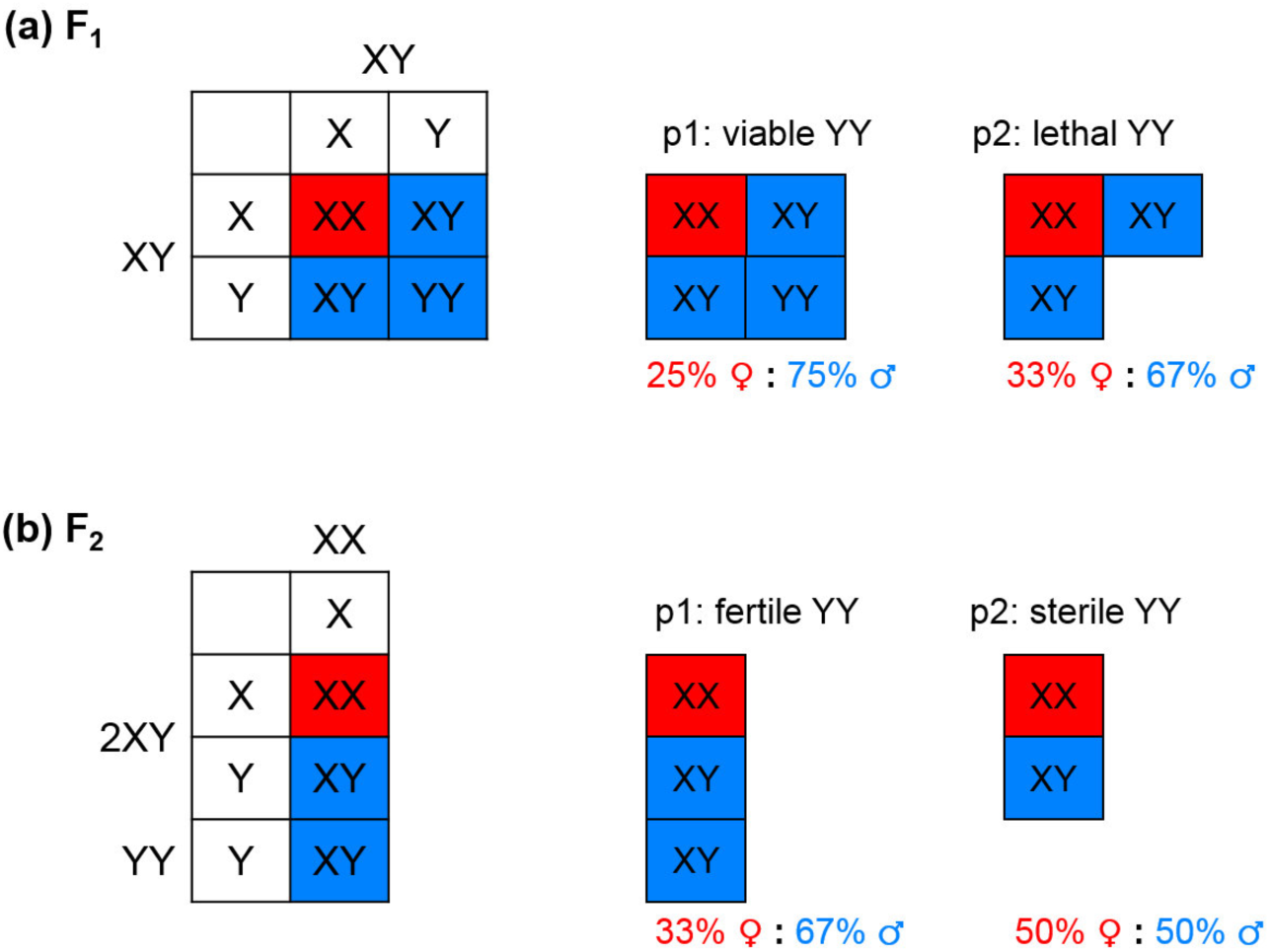
Expected sex ratio under normal and compromised (a) viability and (b) fertility of YY males. **(a)** Viability test. F_1_ progeny generated by crossing XY plants yield a theoretical ratio of 1XX : 2XY : 1YY at the zygote level under the assumption of random mating and no additional mechanism regulating the primary sex ratio. If YY males are as viable as their XY counterparts (p1), the expected F_1_ sex ratio is 0.75, with XY and YY males at a ratio of 2:1. If YY males are completely lethal (p2), the expected sex ratio is then 0.67, with only XY males among the F_1_ progeny. **(b)** Fertility test. F_2_ progeny are generated by random mating between males and females of the F_1_ generation. Crosses between XY and XX yield 50% male and 50% female progeny, while crosses between YY and XX yield only male progeny. Given that the viability test confirmed equal viability of XY and YY males (see Results), the F_1_ generation should have twice the number of XY males as YY males. Random mating these F_1_ males with females should thus yield an F_2_ sex ratio of 0.67 (assuming equal viability of XY and YY males, and thus an equal proportion of XY progeny sired by the two male genotypes). If YY males are completely sterile, however, they will not contribute to the F_2_ generation, and the expected F_2_ sex ratio should thus be 0.50. Red, female. Blue, male.

### Assessment of pollen morphology and anatomy in YY and XY males

We compared the morphology and gross internal structure of pollen grains from air-dried flowers produced by 9 YY and 9 XY male progeny of leaky males from the hormone experiment, using both scanning electron microscopy (SEM) and transmission electron microscopy (TEM). We also included a mixed sample from 5 fresh normal diploid *M. annua* males as a further control. Whole flowers were used for SEM, while anthers were isolated for TEM. Multiple observations were applied to verify the results. Pollen size (polar and equator diameters) was measured on SEM images using ImageJ v1.49 (Schneider *et al.*, 2012).

### Assessment of the relative fertility of YY and XY males

We compared the fertility of YY and XY males by assessing their relative siring success in competition with one another in a common garden, using progeny sex ratios to infer fertility (Figure 1b). Specifically, we grew all progeny produced from the XY crosses from the hormone experiment in a single common garden, i.e. crossing XX sisters with their XY and YY brothers via open pollination. Mature seeds from the female plants were collected and grown to estimate the F_2_ sex ratio (i.e. the proportion of males among the progeny). We reasoned that all seeds sired by YY males would be males, whereas those sired by XY males would have a 50% sex ratio. Given that YY individuals made up 1/3 of the male plants in the F_1_ progeny of XY parents (see Results), we thus expected 67% of the F_2_ progeny to be males if there was equal fertility of YY and XY males (p1 in Figure 1b); in contrast, if YY individuals were completely sterile, all F_2_ progeny should be sired by XY males, with a corresponding 50% sex ratio (p2 in Figure 1b). We applied both Chi-square and likelihood-ratio tests to test for deviation of the observed F_2_ sex ratio from these two extreme scenarios, adjusting for the actual F_1_ XY : YY ratio based on our molecular genotyping described above.

## Results

### Relative viability of YY and XY males

Mating among hormone-feminized males (hormone experiment) or among males with pruning female-flower production (pruning experiment) yielded 939 seeds, of which 278 survived until maturity. Of these, there were 203 male and 75 female progeny (73% males). Recall that random mating among XY males should produce a proportion of 75% male progeny if YY and XY individuals are equally viable (Figure 1a). The observed F_1_ sex-ratio is thus consistent with the scenario of equivalent viability of the two male types (i.e., we failed to reject the null hypothesis p1: X^2^_[1]_ = 0.58, *P* = 0.45). Alternatively, inviability of all YY progeny would have yielded a sex ratio of 67% males; our results are inconsistent with this scenario (p2: X^2^_[1]_ = 5.05, *P* = 0.02). Fully YY viability was ∼18 times more likely than complete YY lethality (Figure S4a). The restriction enzyme genotyping assay confirmed the presence of 57 YY and 106 XY males in our sample of progeny. A Chi-squared test revealed no significant deviation from the expected 2:1 ratio of XY : YY male progeny (X^2^_[1]_ = 0.20, *P* = 0.66).

### Comparisons between YY and XY phenotypes

We found almost no phenotypic differences between YY and XY males, irrespective of whether they had been generated by the pruning experiment or the hormone experiment (Table 1 & 2, Figure S1). Specifically, YY males were identical to normal XY progeny from the hormone experiment in plant height, biomass/height ratio, absolute biomass allocated to different organs (flower, peduncle and vegetative parts), male reproductive allocation (MRA), and the flower/peduncle ratio (*P* > 0.05 for all comparisons; Table 1). Nor were there any differences between the two male genotypes in terms of their principal components from a PCA analysis (Table S1). The results from the pruning experiment were similar, except that YY males had a slightly higher MRA than XY males (N = 53, *P* = 0.025), as well as higher relative reproductive biomass allocation to male flowers (N = 61, *P* = 0.024 for flower/peduncle ratio; Table 2).

### Comparisons of pollen between YY and XY males

Scanning electron microscopy (SEM) revealed clear differences in the morphology and exine ornamentation of pollen produced by YY versus XY males (Figure 2, top row). In general, pollen grains were small, with a polar diameter range of 15 - 28 μm and an equatorial diameter range of 13 - 20 μm (Table S2). Dried fresh pollen grains from XY males were prolate elliptic, tricolpate monads, with a mainly reticulate exine sculpture (Figure 2a). Apertures were invisible due to the constricted colpori. In contrast, dried pollen grains of XY males were elliptic (Figure 2b), with one aperture located in the middle of each slightly infolded lolongate colporus. The ornamentation was granulate and reticulate, with small holes distributed on parts of the pollen wall. Irregular attachments were occasionally observed on the pollen surface. Pollen grains produced by YY males were nearly spheroidal, with mainly granulate ornamentation and more frequent surface presentation of verrucae (Figure 2c).

**Figure 2.**
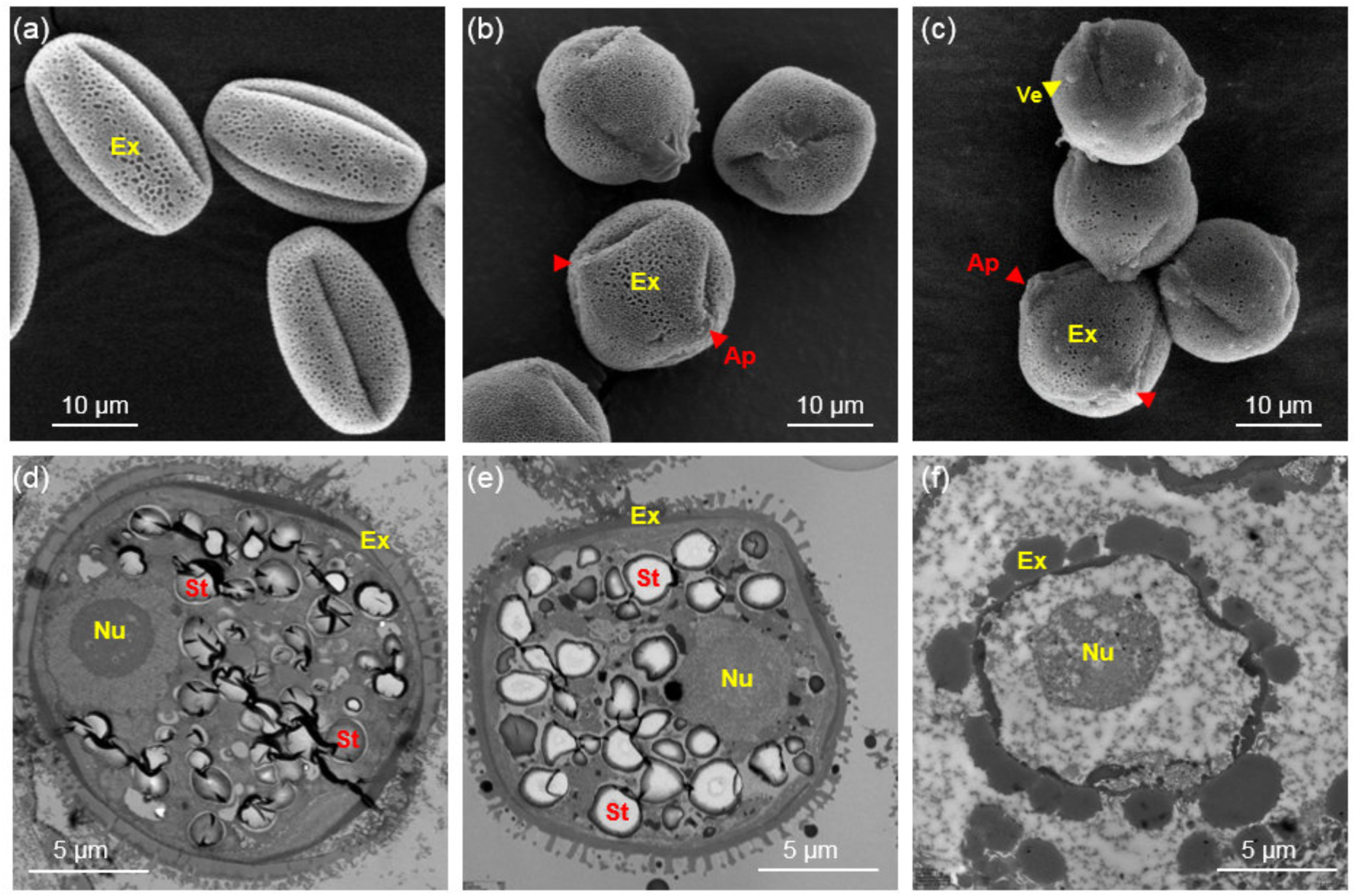
Scanning (top row) and transmission electron microscopy (bottom row) of individual pollen grains. **(a, d)** Fresh pollen from control XY males. **(b, e)** Dried pollen from XY males. **(c, f)** Dried pollen from YY males. Note that both normal-looking **(e)** and abnormal-looking **(f)** samples were observed among specimens of dried pollen from XY and YY plants, but at different proportions (see Supporting Information, Table S2). Ex, exine; Ap, aperture (red arrow); Ve, verruca (yellow arrow); Nu, nucleus; St, starch particle.

Transmission electron microscopy (TEM) revealed more significant differences in the internal ultra-structures between pollen grains produced by XY and YY males (Figure 2, bottom row). Both fresh (Figure 2d) and air-dried pollen grains (Figure 2e) from XY males maintained an internal matrix and had a distinct cell boundary. In contrast, many pollen grains from YY males (Figure 2f) had a thicker exine layer, lacked starch storage within the grain, and presented an indistinct separation of intracellular and intercellular substrates. Other organelles such as the Golgi apparatus and mitochondria were also much less apparent in pollen from YY than from XY individuals. Interestingly, some YY anthers had pollen that resembled that of XY individuals, i.e. the differences just mentioned were not completely categorical at the individual plant level. Perhaps significantly, any differences between pollen grains among YY individuals were always from different anthers (i.e., we did not observe the two types of pollen in the same anther). Among the 23 biological replicates of single anther specimen from YY males, 12 samples showed abnormal pollen characters, while only two out of 11 samples from XY males were different from control XY samples (Table S2).

### Relative fertility and siring success of YY and XY males

Results from mating arrays suggested that YY males were less fertile than XY males. We obtained 192 F_2_ seedlings from seeds produced from mating among the F_1_ progeny of hormone-treated parents. If we assume that the F_1_ population comprised XY and YY males at a 2:1 ratio (along with XX females), the F_2_ generation should comprise 67% males if XY and YY males are equally capable of siring ovules (Figure 1b). The observed F_2_ sex ratio was 56% (107 males, 85 females), representing a significant deficit (p1: X^2^_[1]_ = 9.74, *P* = 0.004), which is compatible with YY males having sired fewer progeny than XY males. Our data cannot reject a scenario in which YY males are completely infertile (p2: X^2^_[1]_ = 2.77, *P* = 0.096). While infertility of YY males was 21.6 times more likely than complete fertility, YY partial fertility is the most likely explanation for our results (Figure S4b).

## Discussion

To evaluate the phenotypic consequences of Y-chromosome evolution, we compared the viability and fertility of YY and XY males of the plant *M. annua*, as well as the morphology, internal anatomy and siring ability of their pollen. YY and XY males were indistinguishable for almost all traits investigated, indicating that genes on the Y chromosome affecting vegetative growth remain functional. We also observed minor potentially positive effects of double Y-chromosome dosage for some reproductive traits of the adult sporophyte. In contrast, pollen produced by YY males appears to be at least partly sterile. Together, our results are consistent with the conclusion, reached on the basis of DNA sequence analysis (Veltsos *et al.*, 2018, 2019), that the non-recombining region of the *M. annua* Y chromosome is only mildly degenerated. They also indicate that the earliest consequences of Y-chromosome evolution in *M. annua* involve fertility.

### No difference in viability or vegetative morphology between YY and XY males

The similar vegetative morphology and viability of YY and XY males of *M. annua* indicate that an X chromosome is not required for effective growth, and that a double dosage of the Y chromosome has little or no effect on the diploid vegetative phenotype. Previous work has documented mild degeneration of the *M. annua* Y chromosome at the sequence level, with a slightly inflated dN/dS ratio for Y-linked genes, implying the accumulation of mildly deleterious mutations, and clear evidence for the pseudogenisation of only one gene of unknown function (Veltsos *et al.*, 2019). It would appear that any Y-chromosome degeneration does not affect sporophyte growth. Given that the Y chromosome accounts for at least an eighth of the genome of *M. annua*, and that a third of the Y chromosome is non-recombining (Veltsos *et al.*, 2018, 2019), it seems unlikely the absence of phenotypic effects of Y-chromosome evolution is due to a paucity of genes on the Y that affect growth and rather points simply to very mild degeneration.

While it is possible that YY individuals suffer from mildly deleterious effects that we were unable to detect, or from effects on traits that we did not measure, the viability and phenotypic similarity between individuals with and without an X chromosome (or with a single or double dose of the Y) are in striking contrast to studies that show substantially poorer performance of YY males compared to their XY counterparts. For instance, Y-chromosome degeneration has led to YY lethality in many animal species (Graves, 2006), as well as in the plants *Silene latifolia* (Westergaard, 1958; Janoušek *et al.*, 1998; Soukupova *et al.*, 2014), and *Carica papaya* (Liu *et al.*, 2004; Ming & Moore, 2007; Yu *et al.*, 2008a). These species probably have older sex chromosomes than *M. annua*, with a longer history of Y-chromosome degeneration. For instance, much of the Y chromosome of *S. latifolia* may have ceased recombining with the X 11 million years ago (Krasovec *et al.*, 2018). In *Carica papaya*, the divergence between X and Y chromosomes has been more recent than in *S. latifolia*, estimated between 0.6 to 2.5 million years (Yu *et al.*, 2008b), but may still be older than that in *M. annua*, in which the non-recombining region is likely less than 1 million years old (Veltsos *et al.*, 2019). The vegetative performance of YY compared to XY individuals of *M. annua* is perhaps similar to that for *Asparagus officinalis* (Harkess *et al.*, 2017) and *Spinacia oleracea* (Wadlington & Ming, 2018), which also have viable YY and nascent sex chromosomes. However, we are unaware of any direct comparisons between YY and XY phenotypes for any plant species other than *M. annua*.

### Difference in reproductive traits and fertility between YY and XY males

In contrast to the purely vegetative traits, we found differences between YY and XY males for certain reproductive traits: YY males had higher reproductive allocation and a greater flower/peduncle ratio than XY males in the pruning experiment. These results are compatible with a scenario in which homozygosity at one or more loci on the Y chromosome results in elevated male flower production, suggesting potential male-beneficial effects on the Y for at least one component of reproductive success. Such ‘super-male’ effects of the YY genotype for both non-reproductive and reproductive traits have been found in some animals, e.g., mating behaviour and siring success in *Oryzias latipes* (Hamilton *et al.*, 1969), but we are not aware of similar reports for any plant species.

Notwithstanding the potentially positive effects of the YY configuration on reproductive traits, our results point clearly to the infertility of YY males. Crossing results indicated that F_2_ progeny were sired more by XY than YY males, and YY males produced a substantial proportion of pollen grains that differed from pollen produced by XY males in external and internal morphological traits. It is noteworthy that pollen development of YY males varied between anthers of the same individual, with some anthers producing only defective pollen and others producing only pollen with normal appearance. It thus appears that the YY configuration compromises the stability of pollen development at the anther level, a suggestion that is coherent with the widely observed temperature sensitivity of male sterility in other plants, where low (e.g. Imin *et al.*, 2004) or high temperatures (e.g. Sakata *et al.*, 2010) render otherwise fertile plants sterile. Although the normal appearance of some of the pollen from YY plants suggests partial fertility, our crossing results do not rule out complete sterility of YY males.

There are two possible explanations for the observed pollen sterility of YY males in *M. annua*. One possibility is that the Y carries a partially penetrant loss-of-function mutation of a gene that is functional on the X, consistent with the mild degeneration of the Y (Veltsos *et al.*, 2018, 2019). If so, our results would imply that Y-chromosome degeneration can have deleterious effects even on an essential male function. Such an implication runs counter to the expectation that the X chromosome should harbor alleles under sexual antagonistic selection that promote female fertility and/or suppress male function, although the X may still contain recessive male-fertility genes (Ellegren & Parsch, 2007). Indeed, the X is known to contain genes associated with spermatogenesis in humans (Ross *et al.*, 2005). An alternative explanation is that the pollen sterility of YY males represents an effect of dosage: a Y-linked gene with male-biased or male-limited expression, of which there are several (Veltsos *et al.*, 2019), may impair pollen development when expressed at a double dose, which is normally not tested by selection. Although we cannot distinguish between these two possibilities without gene expression comparisons between XY, YY and XXY individuals, our results point unambiguously to a divergence in gene function between an X- and Y-linked allele affecting pollen development.

Importantly, the effects of Y-chromosome evolution on pollen development of YY males in *M. annua* differ from the effect of certation, i.e., the poorer performance of Y-than X-bearing pollen tubes, as has been reported for dioecious species of *Rumex* (Stehlik & Barrett, 2005; Stehlik *et al.*, 2008; Sandler *et al.*, 2018). Certation biases the progeny sex ratio towards more daughters, possibly the result of the expression of deleterious mutations in the haploid genome of Y-bearing haploid pollen tubes (Smith, 1963; Lloyd, 1974); such mutations are expected to accumulate through a process of Muller’s ratchet, or because purifying selection is weakened in non-recombining regions by background selection and Hill-Robertson interference (Nei, 1970; Charlesworth, 2002; Vyskot & Hobza, 2004). In contrast, the poor performance of pollen from YY individuals in *M. annua* may be the outcome of a quasi-neutral process, because the vast majority of pollen production in wild populations is by XY individuals (YY individuals are very rare in nature, Cossard and Pannell, 2019), i.e., the effects we have observed would almost never be exposed to purifying selection in the wild. Moreover, in contrast to observations of sex-ratio bias in *Rumex*, the consistently equal sex ratio in dioecious populations of diploid *M. annua* indicates that the Y-bearing pollen grains compete on equal terms with X-bearing pollen in their race to fertilize ovules. We thus have no evidence for effects of the Y chromosome on the haploid gametophytic phase in *M. annua*, be they deleterious and associated with chromosome degeneration, or beneficial and associated with the accumulation of male-beneficial alleles (Scott & Otto, 2017).

### Conclusion

To our knowledge, this is the first report of male sterility associated with the absence of an X chromosome in plants that are otherwise completely viable. It joins evidence from the effects of certation on progeny sex ratios for early phenotypic effects of sex-chromosome evolution (Stehlik & Barrett, 2005; Stehlik *et al.*, 2008), albeit in the context of (diploid) microsporogenesis rather than on (haploid) pollen-tube growth. The cause of the pollen defect is unclear, but the sterility of YY males would seem to suggest either (1) that at least one gene required for the proper development of fertile pollen is compromised on the Y and remains functional on the X, or (2) that a double dose of one or more genes on the Y that have evolved male-biased male-limited gene expression impairs pollen-grain development. Future work comparing patterns of gene expression for such genes between XY and YY males will help to distinguish between these possibilities.

## Acknowledgements

We thank Stephen Wright, Dan Schoen and Marc Johnson for inviting us to contribute to this special issue. JRP thanks Spencer Barrett for many years of stimulating science and mentorship. Yi Huang helped with data collection from the hormone experiment; Yves Cuenot and Dessislava Savova Bianchi helped with the molecular tests; Antonio Mucciolo, Damien De Bellis from the Electron Microscopy Facility at the University of Lausanne provided technical support for electron microscopy. The research was funded by grants from the Swiss National Science Foundation and studentships awarded to XL and JG from the Faculty of Biology and Medicine, University of Lausanne.

## Author contributions

JRP, XL and PV conceived the study. XL, PV and GC conducted the experiments and collected the data. PV and JG developed the sex-specific marker and performed the molecular test. XL performed the electron microscopy. XL and PV analysed the data. XL, PV, JG and JRP wrote the manuscript. All authors approved the final version of the manuscript.

