## Supplemental files for "YY males of the dioecious plant *Mercurialis annua* are fully viable but produce largely infertile pollen"

### Supplementary Figures

**Fig. S1** Comparison in phenotypic traits between XY and YY males of the F1 progeny produced by crossing XY plants in the hormone-induced experiment (‘hormone’) or the pruning-induced experiment (‘pruning’).

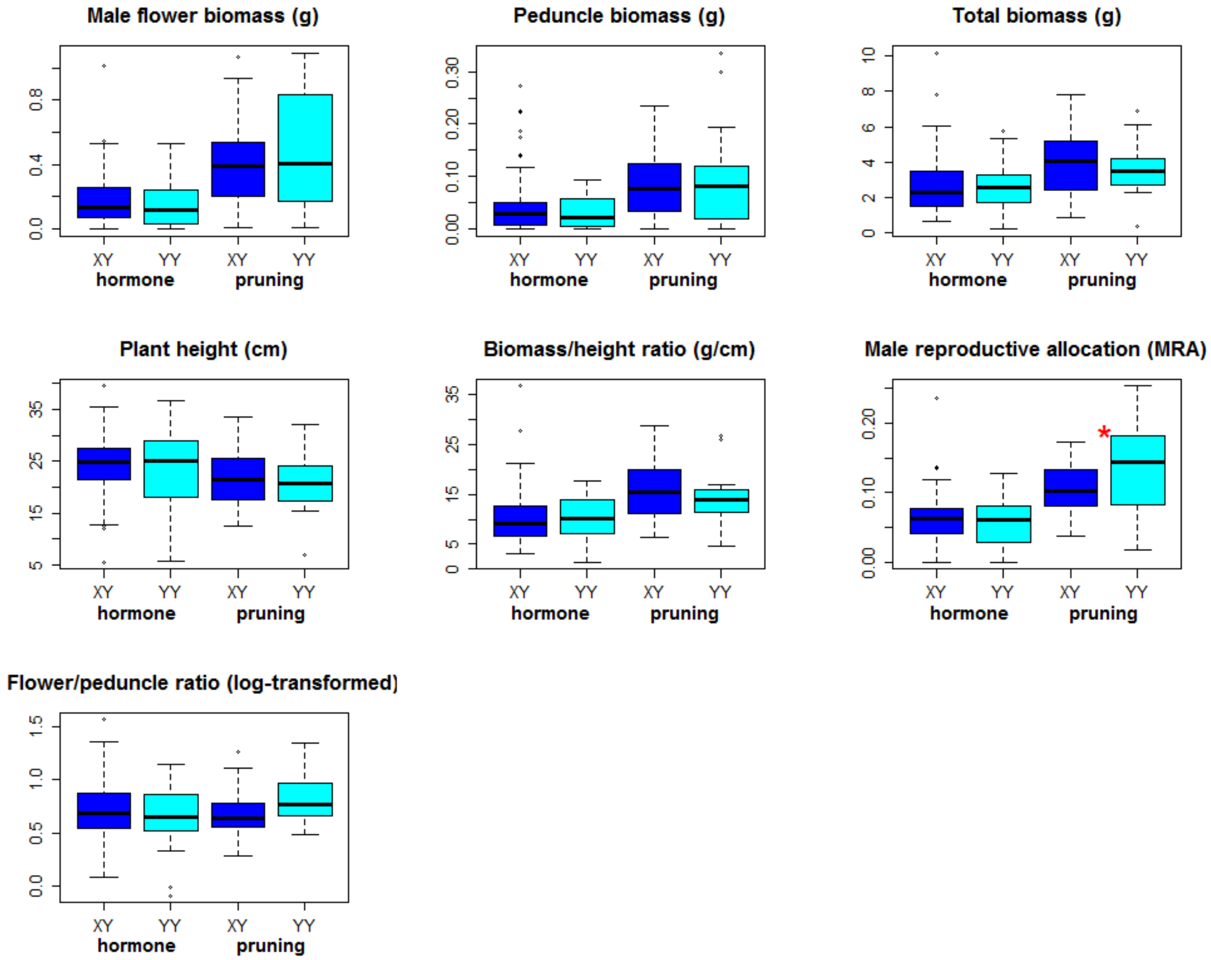


**Fig. S2** Summary of PCA for phenotypic measurements of F1 males from the hormone-induced experiment. Four independent traits are included for the analysis after normalisation: male flower biomass (flower_biomass), peduncle biomass (peduncle_biomass), total biomass (dry_biomass) and plant height (height).

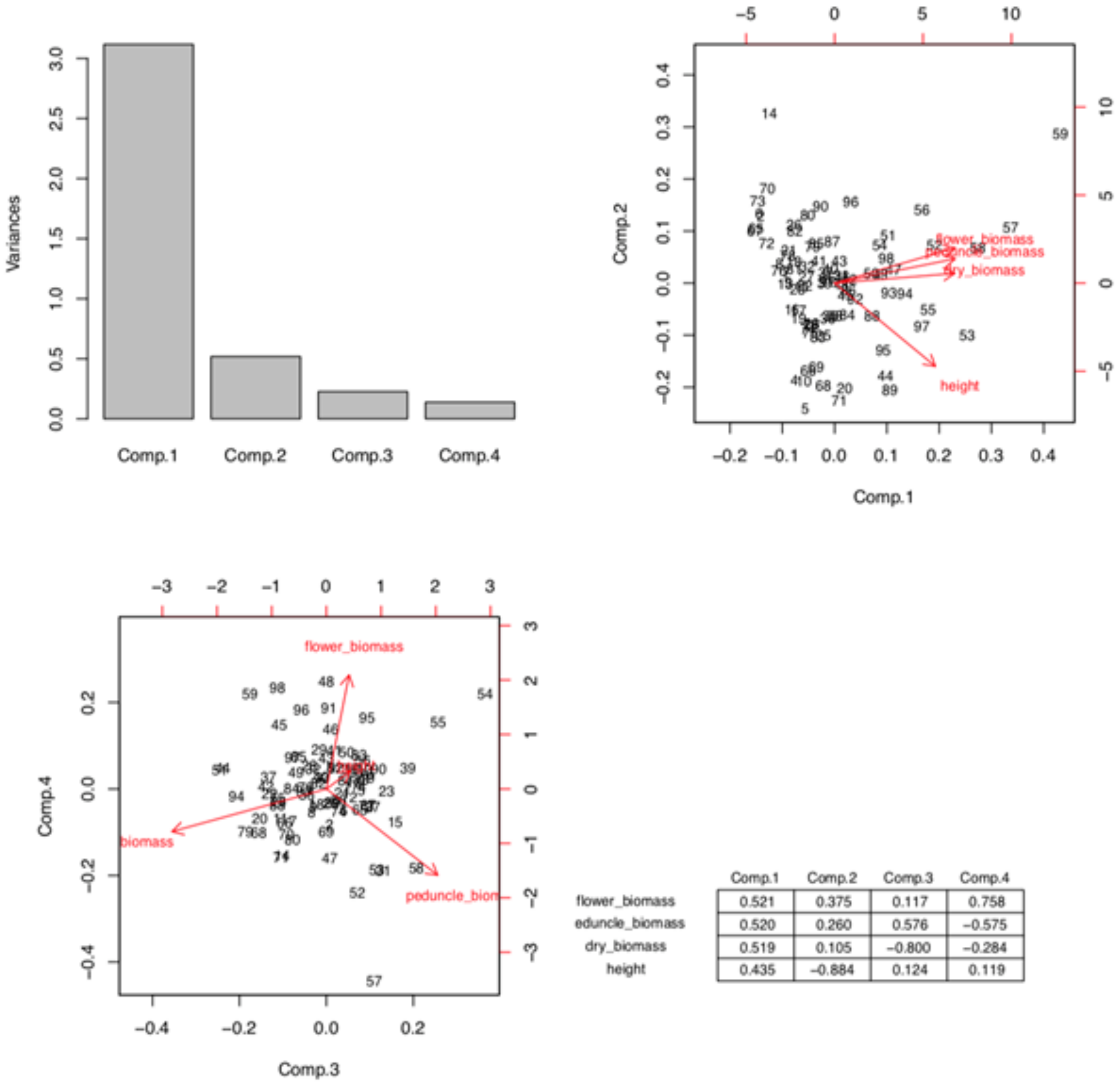


| | Comp.1 | Comp.2 | Comp.3 | Comp.4 |
|---|---|---|---|---|
| flower_biomass | 0.521 | 0.375 | 0.117 | 0.758 |
| eduncle_biomass | 0.520 | 0.260 | 0.576 | −0.575 |
| dry_biomass | 0.519 | 0.105 | −0.800 | −0.284 |
| height | 0.435 | −0.884 | 0.124 | 0.119 |

**Fig. S3** Summary of PCA for phenotypic measurements of F1 males from the pruning-induced experiment.

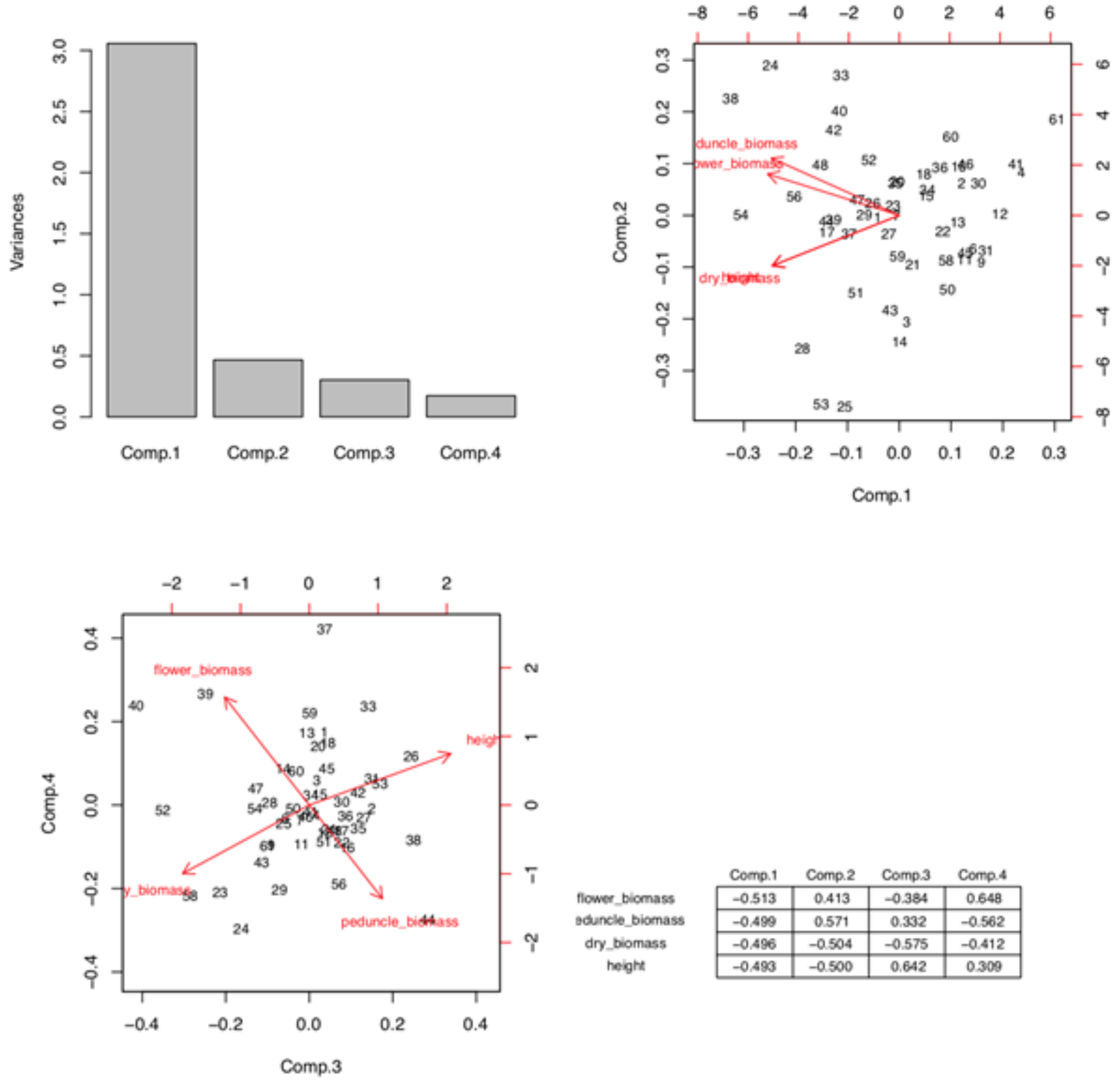


| | Comp.1 | Comp.2 | Comp.3 | Comp.4 |
|---|---|---|---|---|
| flower_biomass | −0.513 | 0.413 | −0.384 | 0.648 |
| eduncle_biomass | −0.499 | 0.571 | 0.332 | −0.562 |
| dry_biomass | −0.496 | −0.504 | −0.575 | −0.412 |
| height | −0.493 | −0.500 | 0.642 | 0.309 |

**Fig. S4** Likelihood ratio tests for progeny sex ratios. **(a)** Likelihood ratio test on the proportion of males among $F_1$ progeny. p1, equal viability between XY and YY males, resulting in an expected $F_1$ sex ratio of 0.75; p2, complete YY lethality, resulting in an expected $F_1$ sex ratio of 0.67. **(b)** Likelihood ratio test on the proportion of males among $F_2$ progeny. p1, equal fertility between XY and YY males, resulting in an expected $F_2$ sex ratio of 0.67; p2, complete YY sterility, resulting in an expected $F_2$ sex ratio of 0.5.

**(a)**

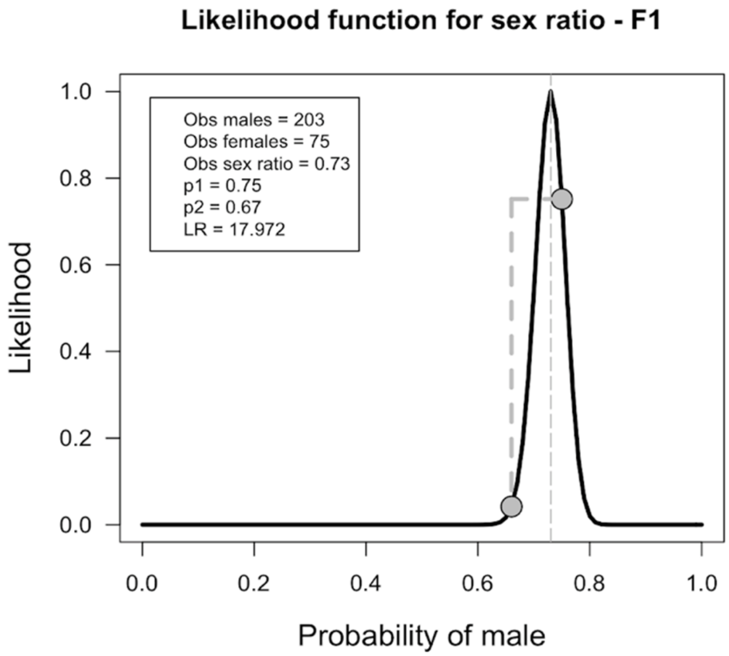


**(b)**

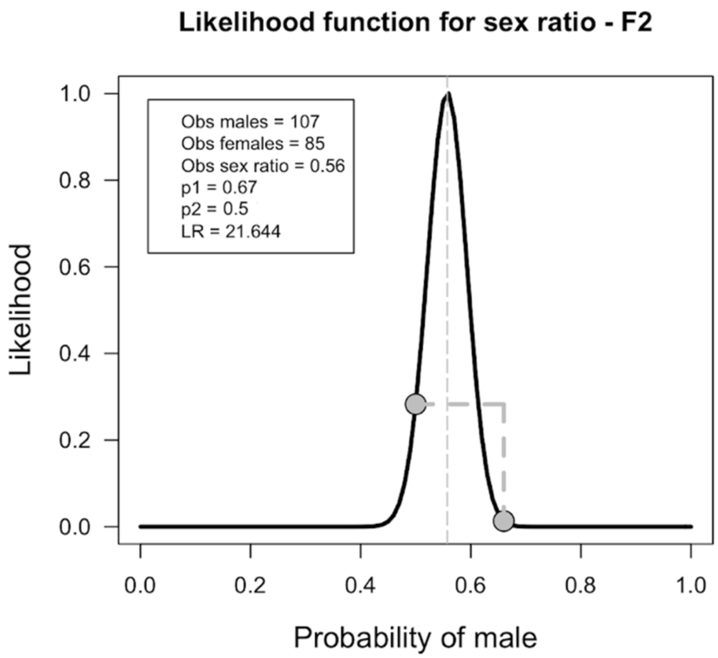

**Fig. S5** Top: Illustration of the cutting position of the restriction enzymes that only cut the X or the Y allele. Bottom: Illustration of the restriction enzyme assay to distinguish XY and YY individuals.

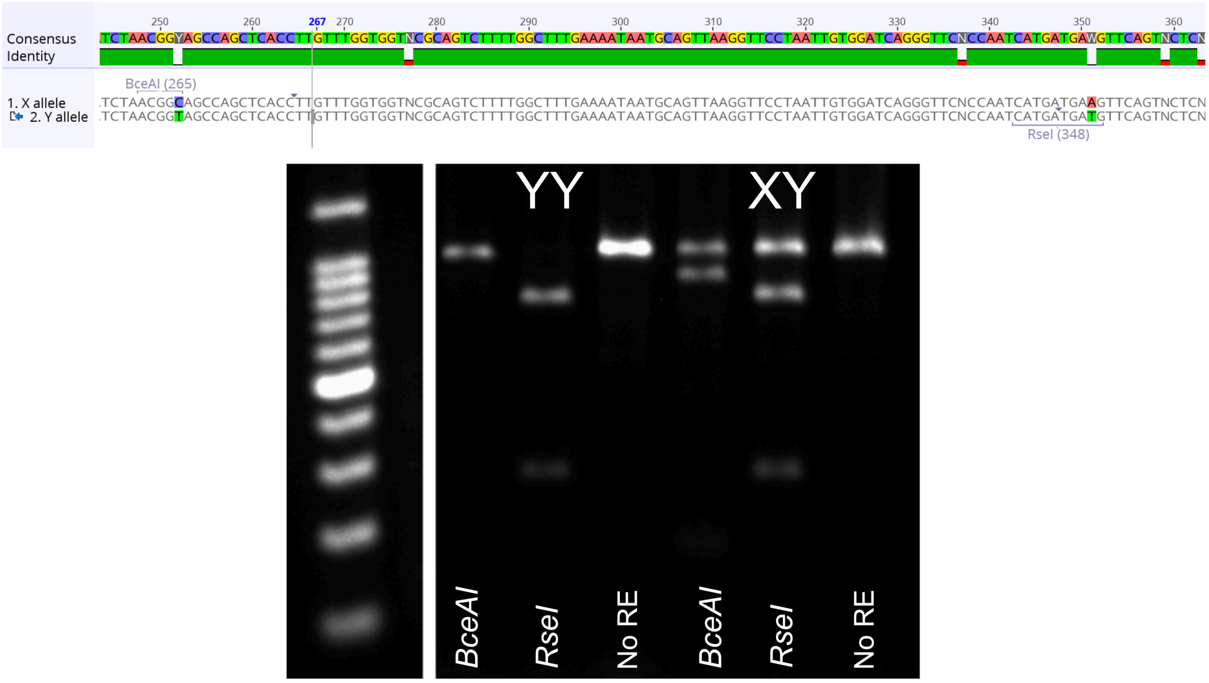

### Supplementary Tables

**Table S1** Summary of generalised linear mixed model on binomial genotype (XY versus YY) explained by the first 4 principal components of normalised independent phenotypic measurements. Left: male progeny from the hormone-induced experiment, with sampler and sampling date as random variables. Right: male progeny from the pruning-induced experiment, with family as a random variable.

| | Dependent Variables | |
|---|---|---|
| Predictors | genotype | |
| | *Odds ratio (CI)* | *p* |
| **Fixed Parts** | | |
| (Intercept) | 0.60 (0.38 – 0.94) | **0.026** |
| Comp.1 | 0.84 (0.62 – 1.12) | 0.239 |
| Comp.2 | 0.91 (0.49 – 1.69) | 0.772 |
| Comp.3 | 0.43 (0.16 – 1.20) | 0.106 |
| Comp.4 | 1.31 (0.33 – 5.19) | 0.703 |
| **Random Parts** | | |
| $\tau_{00, date}$ | 0.000 | |
| $\tau_{00, sampler}$ | 0.000 | |
| $N_{date}$ | 3 | |
| $N_{sampler}$ | 2 | |
| $ICC_{date}$ | 0.000 | |
| $ICC_{sampler}$ | 0.000 | |
| Observations | 86 | |
| AIC | 124.292 | |
| Deviance | 110.292 | |

| | Dependent Variables | |
|---|---|---|
| Predictors | genotype | |
| | *Odds ratio (CI)* | *p* |
| **Fixed Parts** | | |
| (Intercept) | 0.36 (0.18 – 0.72) | **0.004** |
| Comp.1 | 1.06 (0.74 – 1.52) | 0.744 |
| Comp.2 | 3.44 (1.06 – 11.21) | **0.040** |
| Comp.3 | 0.85 (0.26 – 2.76) | 0.783 |
| Comp.4 | 4.19 (0.89 – 19.74) | 0.070 |
| **Random Parts** | | |
| $\tau_{00, family}$ | 0.000 | |
| $N_{family}$ | 19 | |
| $ICC_{family}$ | 0.000 | |
| Observations | 53 | |
| AIC | 68.030 | |
| Deviance | 56.030 | |

**Table S2** Pollen observation under electron microscopy

| Sample type | $N_{individuals}$ | Pollen size under SEM (μm) | | | TEM | |
|---|---|---|---|---|---|---|
| | | $N_{pollen}$ | Polar diameter (mean ± SE) | Equator diameter (mean ± SE) | $N_{normal}$ | $N_{abnormal}$ |
| Dried pollen from YY males | 9 | 33 | 21.96 ± 3.64 | 17.52 ± 0.99 | 11 | 12 |
| Dried pollen from XY males | 9 | 36 | 22.70 ± 1.54 | 17.50 ± 1.18 | 9 | 2 |
| Fresh pollen from XY males (control) | 5 | 30 | 26.64 ± 1.16 | 15.23 ± 0.84 | 8 | 0 |

*SEM, scanning electron microscopy; TEM, transmission electron microscopy.

**Methods S1** Identification between XY and YY males with a restriction enzyme-based assay based on sex-specific SNPs

We used previously published genome capture sequencing data to identify one exon with male-specific SNPs across 10 populations spanning most of the species range (2 males and 2 females in each, see González-Martínez *et al.*, 2017 for details). We confirmed the presence of this SNP by direct Sanger sequencing of PCR products that amplify in both sexes in individuals from Barcelona (SP), Antalya (TR), Jerusalem (ISR), Southampton (UK), Trabzon (TR) and Volos (GR). The Sanger sequences (Fig. **S5**, top) were aligned using MAFFT (Katoh & Standley, 2013) in Geneious v9.1.5 (https://www.geneious.com). PCR amplification with primers 1911F (CGCTGTGACTGATCCTGGAA) and 1911R (CTCTTGAGATTGGCCTGCCA) at 55°C annealing temperature, 1 min extension at 72°C and 30 PCR cycles and resulted in a 727 bp product. The product contained two SNPs and could be reliably digested with restriction enzymes *BceAI* (X-specific) and *RseI* (Y-specific) separately, to result in an assay that distinguishes XY and YY individuals (Fig. **S5**, bottom).

**González-Martínez SC, Ridout K, Pannell JR**. **2017**. Range expansion compromises adaptive evolution in an outcrossing plant. *Current Biology* **27**: 2544–2551.e4.

**Katoh K, Standley DM**. **2013**. MAFFT multiple sequence alignment software version 7: improvements in performance and usability. *Molecular Biology and Evolution* **30**: 772–780.
